# Blood oxygenation-level dependent cerebrovascular reactivity imaging as strategy to monitor CSF-hemoglobin toxicity

**DOI:** 10.1101/2022.04.05.487168

**Authors:** Bart R. Thomson, Henning Richter, Kevin Akeret, Raphael M. Buzzi, Vania Anagnostakou, Christiaan H. B. van Niftrik, Nina Schwendinger, Zsolt Kulcsar, Peter W. Kronen, Luca Regli, Jorn Fierstra, Dominik J. Schaer, Michael Hugelshofer

## Abstract

**Purpose:** Cell-free hemoglobin in the cerebrospinal fluid (CSF-Hb) may be one of the main drivers of secondary brain injury after aneurysmal subarachnoid hemorrhage. Haptoglobin scavenging of CSF-Hb has been shown to mitigate cerebrovascular disruption. Using digital subtraction angiography (DSA) and blood oxygenation-level dependent cerebrovascular reactivity imaging (BOLD-CVR) the aim was to assess the acute toxic effect of CSF-Hb on cerebral blood flow and autoregulation, as well as to test the protective effects of haptoglobin.

**Methods:** DSA imaging was performed in eight anesthetized and ventilated sheep (mean weight: 80.4 kg) at baseline, 15, 30, 45 and 60 minutes after infusion of hemoglobin (Hb) or co-infusion with haptoglobin (Hb:Haptoglobin) into the left lateral ventricle. Additionally, 10 ventilated sheep (mean weight: 79.8 kg) underwent BOLD-CVR imaging to assess the cerebrovascular reserve capacity.

**Results:** DSA imaging did not show a difference in mean transit time or cerebral blood flow. Wholebrain BOLD-CVR compared to baseline decreased more in the Hb group after 15 minutes (Hb vs Hb:Haptoglobin: −0.03 ±0.01 vs −0.01 ±0.02) and remained diminished compared to Hb:Haptoglobin group after 30 minutes (Hb vs Hb:Haptoglobin: −0.03 ±0.01 vs 0.0 ±0.01), 45 minutes (Hb vs Hb:Haptoglobin: −0.03 ±0.01 vs 0.01 ±0.02) and 60 minutes (Hb vs Hb:Haptoglobin: −0.03 ±0.02 vs 0.01 ±0.01).

**Conclusion:** It is demonstrated that CSF-Hb toxicity leads to rapid cerebrovascular reactivity impairment, which is blunted by haptoglobin co-infusion. BOLD-CVR may therefore be further evaluated as a monitoring strategy for CSF-Hb toxicity after aSAH.

## INTRODUCTION

Aneurysmal subarachnoid hemorrhage (aSAH) is a severe form of intracranial bleeding caused by the rupture of an intracranial aneurysm^1^, which accounts for 5-10% of all strokes^2^. In addition to the early brain injury occurring within the first 72 hours after SAH^3^, patient outcome is mainly determined by secondary brain injury (SAH-SBI), which occurs between 3-14 days after the initial bleeding^4^. Three main components of SAH-SBI are identified: 1. angiographic vasospasm in large cerebral arteries (aVSP); 2. radiologically diagnosed delayed cerebral ischemia; and 3. clinically evident delayed ischemic neurologic deficits. SAH-SBI is a complex process and recent evidence indicates that one of the main drivers may be the acute toxicity of cell-free hemoglobin in the cerebrospinal fluid (CSF-Hb)^5^.

The pathophysiology is believed to involve the gradual lysis of erythrocytes in the subarachnoid space, thereby releasing large amounts of Hb^5,6^, causing direct oxidative tissue injury^7,8^ and interaction with nitric oxide (NO) signaling in the arterial vessel wall, resulting in cerebrovascular disruption^9,10^. Recent findings from our group^10^ and others^11^ suggest that scavenging of CSF-Hb by haptoglobin mitigates these negative effects by preventing CSF-Hb tissue penetration, thus maintaining NO signaling. However, it remains unknown whether CSF-Hb toxicity induced cerebrovascular disruption primarily occurs on the level of major cerebral arteries or if toxic effects on microcirculation^11, 12^ significantly disturb the autoregulatory control of cerebral perfusion. To answer this question, a translational sheep model with direct intraventricular injection of cell-free Hb or co-infusion of Hb with haptoglobin was used to observe macrovascular effects using digital subtraction angiography (DSA)^13^. Additionally, blood oxygenation-level dependent cerebrovascular reactivity (BOLD-CVR) imaging was performed to assess effects of CSF-Hb on the cerebrovascular autoregulatory reserve capacity on a microvascular level^14,15^. Imaging strategies, which allow for detection of acute CSF-Hb effects in aSAH patients may provide important guidance for clinical studies targeting these toxicity pathways.

## METHODS

### General

This study was conducted according to the Swiss Animal Welfare Act (TschG, 2005) and the Swiss Animal Welfare Ordinance (TSchV, 2008) received ethical approval from the Swiss Federal Veterinary Office Zurich (animal license no. ZH 234/17). The authors complied with the ARRIVE guidelines. A total of 20 female Swiss alpine sheep, aged 2-4 years and obtained from the Staffelegghof (see supplemental material) were used during this study. Eight sheep were used during the DSA experiment (n= 4 Hb; n=4 Hb:Haptoglobin) and twelve during the BOLD-CVR experiment (n=6 Hb; n=6 Hb:Haptoglobin, Figure 1B). Prior to the experiments the animals underwent a health check (clinical and blood examination) by a veterinarian and were randomly assigned to treatment groups. All involved personnel were blinded for group allocation until analysis of data.

**Figure 1.**
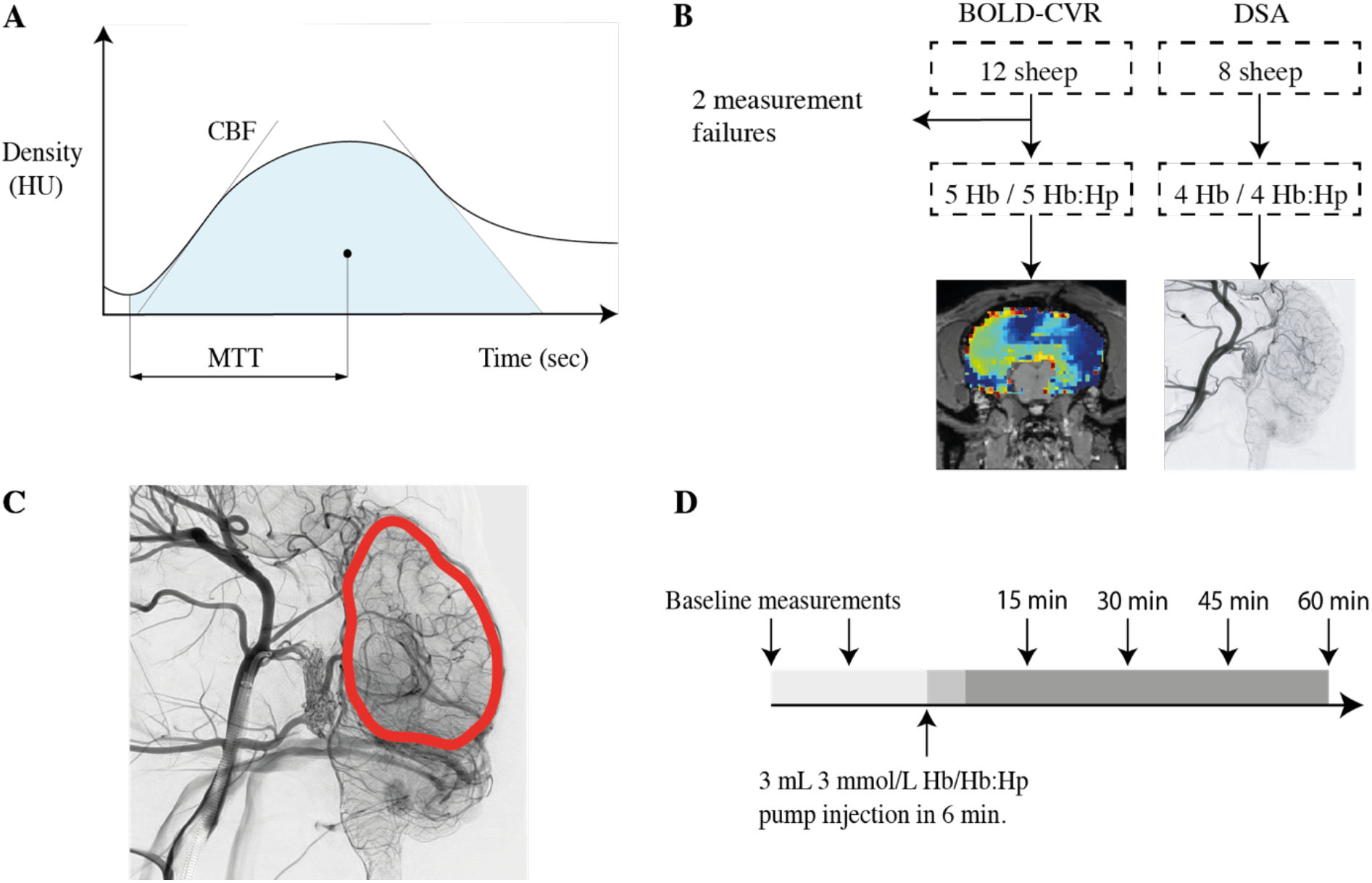
Density curve analysis and DSA region of interest. (**A**) DSA-derived flow parameters MTT and CBF (**B**) Illustration of the experimental workflow (**C**) Full brain region of interest (indicated in red) used for computation of DSA parameters (**D**) Scanning protocol, two baseline acquisitions prior to substance infusion, followed by four acquisitions post substance infusion.

In both experiments, the order of substance infusion per animal was spread homogeneously to minimize potential confounding influences. In each experiment, the parameters of two scans prior to substance infusion were averaged and are presented as the baseline. In the BOLD-CVR experiment, two animals were excluded due to substance infusion failure and due to a physiologically impossible signal response in the baseline scans, identifying the baseline scans as corrupt. Under general anesthesia, surgical navigation was used to instrument the ventilated sheep with a left frontal external ventricular drain (EVD, DePuys Synthes). The model is further described in detail in the appendix. A PHD Ultra syringe pump (Harvard Apparatus) was connected to the EVD through which an excess (3 mL, 3 mmol/L) of Hb or Hb:Haptoglobin was infused within 6 minutes. In both experiments half of the group was infused with Hb, and the other half with Hb:Haptoglobin complex (Figure 1D).

### Hemoglobin and Hemoglobin-Haptoglobin complexes

Hb was purified from sheep blood as previously described^16^. Hb concentrations were determined by spectrophotometry as described and are given as molar concentrations of total heme (1M Hb tetramer is, therefore, equivalent to 4M heme)^9^. For all Hb used in these studies, the fraction of ferrous oxyHb (HbFe2+O2) was always greater than 98% as determined by spectrophotometry. Haptoglobin from human plasma (phenotype 1-1) was obtained from CSL Behring.

### DSA imaging

Allura Clarity angiography suite (Philips Healthcare, Best, The Netherlands) was used to perform DSA with an angiographic 5F catheter (Cordis) placed into the largest anastomotic branch of the right maxillary artery supplying the extradural rete mirabile and the internal carotid artery. Standardized pump injections of contrast media were performed to ensure comparability between pre- and post-substance infusion images. Prior to infusion of the respective substance, two DSA acquisitions were performed, followed by four acquisitions at respectively 15, 30, 45 and 60 minutes after the start of substance infusion.

### DSA data processing

Prior to post-processing the whole cerebrum was segmented (Figure 1C) in the 2D perfusion image. Sequentially, two parameters (Figure 1A) were extracted and analyzed with a customized algorithm in Python. Mean transit time (MTT) was defined as the time from arrival of the contrast fluid till the center of mass of the density curve. Cerebral blood flow (CBF) was derived as cerebral blood volume (CBV) / MTT, where CBV is defined as the area under the contrast density curve.

### BOLD imaging

MR images were acquired on a 3 Tesla MR unit (Philips Ingenia) with a 32-channel head coil. The scanning protocol consisted of a T1-weighted sequence, two 2D EPI BOLD fMRI sequences, followed by a T2-weighted sequence during which infusion of the respective substance took place and ultimately four 2D EPI BOLD fMRI sequences at respectively 15, 30, 45 and 60 minutes after the onset of substance infusion. In one animal in the Hb:Haptoglobin group, scan acquisition at the 60 minutes time point could not be completed due to time constraints. In all acquisitions whole-brain volumes were acquired with a 2×2×2 mm3 voxel size. Additional fMRI parameters were an acquisition matrix of 112×112×33 slices with ascending interleaved acquisition without slice gap, repetition time (TR)/ echo time (TE) 1896/17 ms, a 75-degree flip angle with a bandwidth of 1900 Hz/Px. During acquisition PetCO2 was maintained for 120 s at normocapnia, followed by an abrupt hypercapnic step increase of 15 mmHg for 240 s, before returning to normocapnia for 360 s. Normocapnia depended on the resting PetCO2 of the individual animal, as these were found to range in between individual sheep. The hypercapnic step was achieved by an automated gas blender that adjusts the gas flow and composition to a sequential gas delivery breathing circuit (RepirAct, Thornhill Research Institute, Toronto, Canada) to control the end-tidal partial pressures of PetCO2 and oxygen (PetO2), as described in ^15,17^.

### BOLD-CVR data processing

All acquired imaging was preprocessed using Statistical Parametric Mapping 12 (SPM 12, Wellcome Trust Centre for Neuroimaging, Institute of Neurology, University College London, UK). All temporal BOLD volumes were realigned to the mean of their series, followed by a registration of the anatomical volumes to the same mean image. Ultimately, the functional images were smoothed with a Gaussian kernel of 8mm full width at half maximum. Consecutively, the anatomical acquisitions were used to manually delineate the cerebrum, using 3DSlicer 4.11.0^18^.

Further temporal processing was performed in Python, where linear detrending of the data was followed by, like Duffin et al. (2015), a low band-pass filter with a filter cut-off of 0.125 Hz and lowess smoothing^19^ using 16 dynamics (6%). Ultimately, the PetCO2 course was resampled to match the TR of the BOLD data. CVR calculations were performed after conclusion of the experiments based on a standardized method presented by van Nifrik et al.^20^.

## RESULTS

In the DSA experiment, MTT increases slightly in both groups (Figure 2A & 2B), and CBF (in mL/100g/min) shows a slight decrease (Figure 2C & 2D). However, no apparent differences are observed between the groups following their respective injection. When imaged with BOLD-fMRI, all animals injected with Hb demonstrate a decline in CVR as early as 15 minutes post infusion, after which it stabilizes on this reduced level (Figure 2E and 2F). In contrast, the Hb:Haptoglobin group demonstrates an increase in CVR. Figure 3 presents the decrease in CVR with respect to the baseline following Hb infusion, whereas an increase in CVR is observed after injection of Hb:Haptoglobin. Here, the overall decrease in CVR appears more apparent in cortical regions in proximity of the CSF space, however, no clear difference could be quantified between gray and white matter.

**Figure 2.**
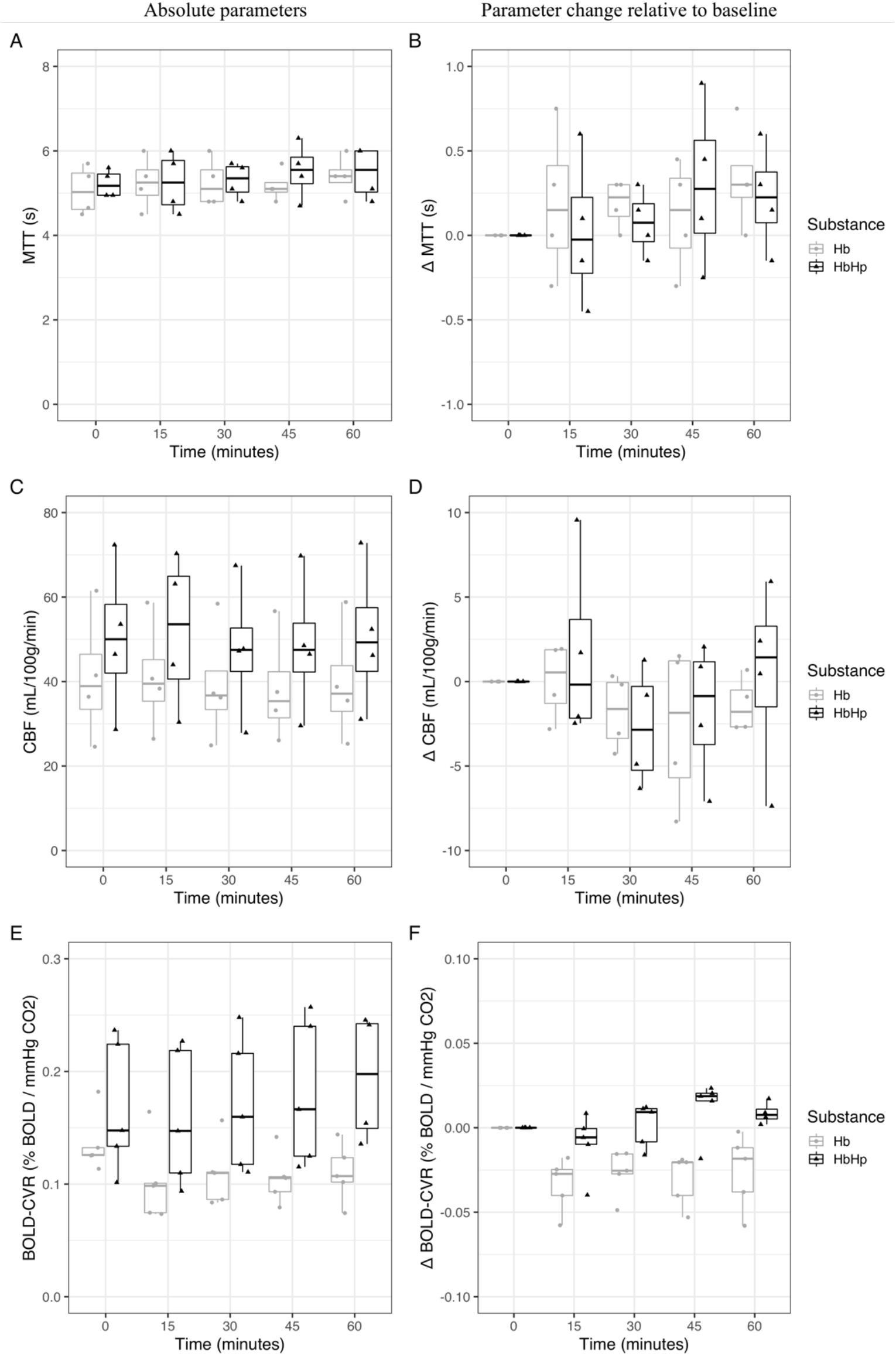
Temporal dynamics of cerebrovascular parameters after intracerebroventricular Hb or Hb:Haptoglobin. Absolute temporal profiles of DSA derived parameters (**A**) MTT, (**C**) CBF and (**E**) BOLD-CVR. Changes with respect to baseline of DSA derived parameters (**B**) MTT (**D**), CBF and (**E**) BOLD-CVR. Hb is indicated in gray and Hb:Haptoglobin (Hb:Hp) in black.

**Figure 3.**
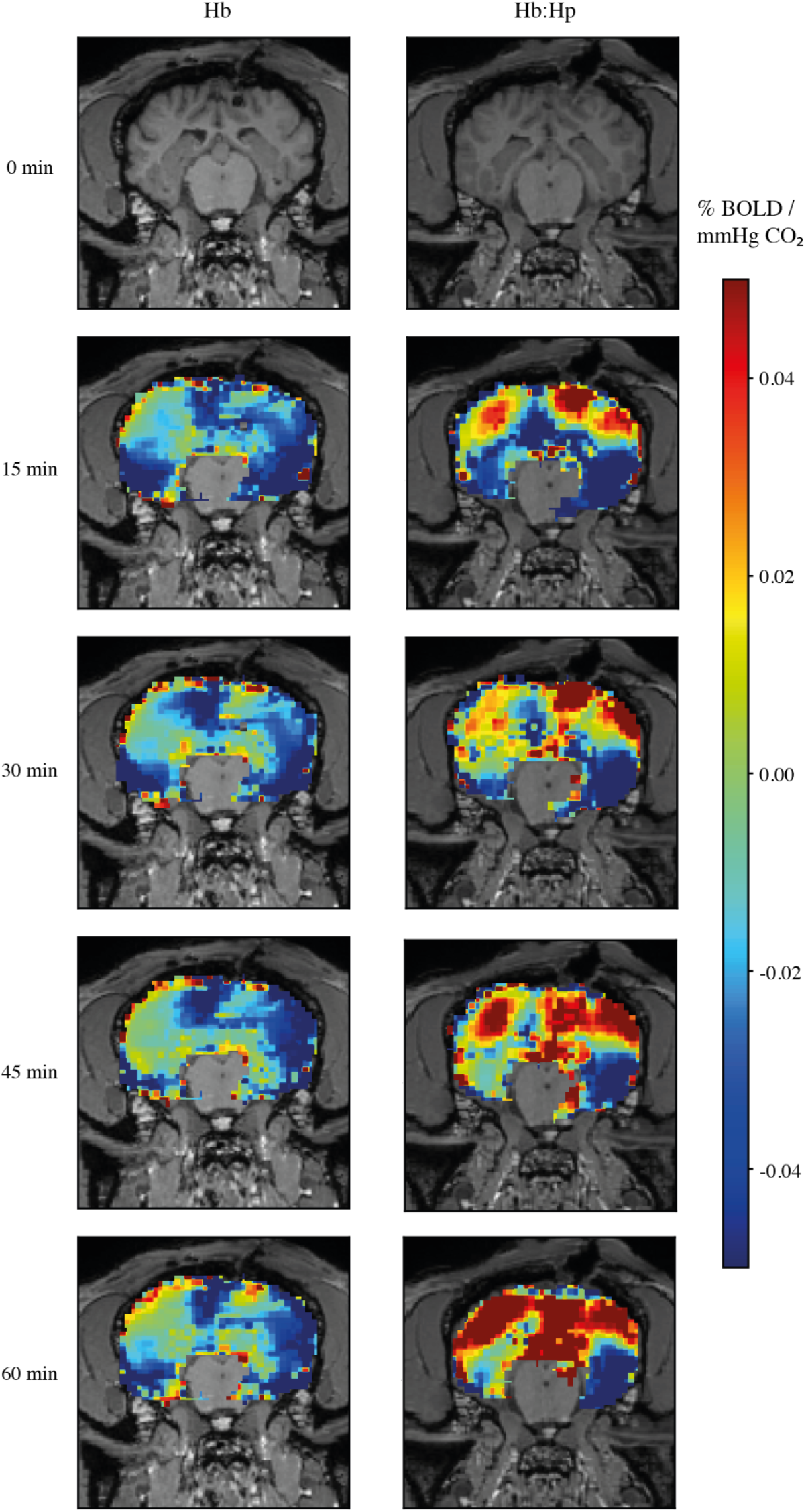
Spatio-temporal dynamics of BOLD-CVR after intracerebroventricular Hb or Hb:Haptoglobin. Spatio-temporal CVR group mean changes after baseline following infusion of 3 mL 3mmol/L Hb (left column) and Hb:Haptoglobin (Hb:Hp, right column) into the CSF space. Colorbar indicates CVR change with respect to baseline.

## DISCUSSION

Our study demonstrates that BOLD-CVR imaging is able to detect early-onset toxicity of CSF-Hb, which can be prevented by haptoglobin co-infusion. Contrarily, the functional DSA derived parameters MTT and CBF were not sensitive enough to detect the difference between Hb or Hb:Haptoglobin infusion. These findings strengthen the rationale for BOLD-CVR monitoring as a strategy to detect CSF-Hb toxicity and haptoglobin treatment effects after aSAH.

In our model, CSF-Hb induced CVR impairment can be detected as early as 15 minutes after Hb infusion into the CSF and therefore precedes Hb-induced macrovascular constriction observed in previous studies^10^. The decreased cerebrovascular response lasts up to 1 hour after infusion of Hb, when compared to baseline CVR. Interestingly, after co-infusion of Hb with haptoglobin we found an increase of CVR over time, when compared to baseline. Upon qualitative assessment these protective effects seem larger in the cortical regions in proximity to the subarachnoid space. Physiologically this could be explained by the cortico-ventricular direction of CSF flow along the perivascular spaces, diluting Hb with highest concentrations found in juxtacortical regions^21^.

In contrast to BOLD-CVR, the DSA derived parameters MTT and CBF were not sensitive enough to detect variations between Hb or Hb:Haptoglobin infusion. These differences are explained by the microvascular changes that are better reflected by BOLD-CVR versus the macrovascular changes detectable by DSA. This could partially explain the clinical observation of occurrence of SAH-SBI in absence of angiographic vasospasm^22^, and presence of vasospasm when SAH-SBI is absent^23^. A previous clinical study^5^ showed strong evidence for a positive association between the occurrence of SAH-SBI and daily measured CSF-Hb concentrations. Future research should be scoped towards relating quantitative BOLD-CVR measurements in aSAH patients to CSF-Hb concentration and the clinical presentation of SAH-SBI. In this regard, the BOLD-CVR results that we present, suggest a possible role for BOLD-CVR as a powerful clinical imaging strategy to guide therapeutic interventions to prevent CSF-Hb toxicity in the brain.

Due to the CBF surrogacy that is reflected by BOLD-CVR, we argue that already after 15 minutes, CSF-Hb affects CBF on a microvascular level. Other groups suggest that there is a link between impaired CVR and the risk of ischemic events^24–25^. Hence, the observed effect following unbound Hb infusion might contribute to the occurrence of delayed cerebral ischemia and delayed ischemic neurologic deficits following aSAH. Moreover, haptoglobin co-infusion seems to prevent these microvascular changes. These findings further support the therapeutic concept of intracerebroventricular haptoglobin administration as strategy to target CSF-Hb toxicity after aSAH^5^, which is planned to be translated into clinical studies in the near future.

The sensitivity of BOLD-CVR imaging to detect CSF-Hb effects on cerebral blood flow regulation may be of high relevance in a clinical context. Since it appears that the BOLD-CVR signal changes forego the manifestation of delayed cerebral ischemia or delayed ischemic neurologic deficits, early detection of CSF-Hb toxicity may allow for targeted therapeutic interventions with the aim to improve microvascular perfusion. Consequently, the demonstrated sensitivity of BOLD-CVR harbors the potential to lower morbidity and mortality associated with SAH-SBI.

Our study contains several limitations. Firstly, our model was designed to show the temporospatial vascular effects of CSF-Hb on CBF regulation. BOLD-CVR is assumed to be a surrogate marker for CBF based on the strong correlation between CBF measured by arterial spin labeling and the BOLD signal^26^. However, the BOLD signal has a complex dependency upon hematocrit, cerebral blood volume and cerebral metabolic rate and is not only dependent on cerebral blood flow^27^. Our experimental setup did not allow for appropriate anatomical quantification of Hb-induced macrovascular constrictions in the BOLD-CVR experiments. The absence of simultaneous anatomical imaging makes it impossible to directly quantify the physiological contribution of macrovasospasm to the decrease in CVR. However, the absence of macrovascular changes in the DSA group hint towards absence of a macrovascular contribution in the reduction of CVR. Secondly, baseline CVR differences are present between the Hb and Hb:Haptoglobin group, which is most likely due to the small sample size, however, still within a range observed within healthy human subjects^28^. This small sample size limited quantitative statistical testing. However, as the clinically relevant differences in CVR were expected to be small, a sample size with adequate power for statistical analysis in these large animal experiments has been judged inappropriate due to animal welfare considerations. Lastly, a sheep model was chosen due to the similar cortical architecture and size when compared to humans^29^, allowing for adequate imaging. To realize a stable model, anesthesia was maintained with isoflurane, which in itself does not affect CSF production, however dosage has been shown to change the reabsorption^30^. Regardless of the effects, the situations in our experiments were reproducible and comparable.

### Conclusions

In conclusion, we have demonstrated that BOLD-CVR is able to quantify microvascular effects of CSF-Hb, prior to macrovascular constriction. This sensitivity allows for earlier adaptive measures and guide therapeutic interventions such as scavenging of CSF-Hb with intracerebroventricular application of haptoglobin. Contrarily, the DSA derived parameters MTT and CBF were not sensitive enough to discriminate between Hb and Hb:Haptoglobin co-infusion. Our results contribute to establishing BOLD-CVR as an imaging modality to detect vascular changes after aSAH with a high sensitivity and strengthen the rationale for novel treatment strategies targeting Hb-toxicity after aSAH.

## Data availability statement

Data and code can be made available upon reasonable request via author correspondence.

