## Supplemental Methods for "Blood oxygenation-level dependent cerebrovascular reactivity imaging as strategy to monitor CSF-hemoglobin toxicity"

**Animals and housing**

Female Swiss alpine sheep (Staffelegghof, Küttigen bei Aarau, Switzerland) with a median weight of 80.5 kg (weight range: 69.5 – 89.0 kg) and an age between 2-4 years were transferred to the Veterinary Hospital of Zurich and allowed to acclimatize to their new environment for at least 7 days. A veterinarian performed standardized screening by means of physical examination and blood testing during this period. Prior to surgery the animals were fasted with free access to water for 18-24 hours.

**Anesthesia**

30 minutes prior to anesthesia, the attending veterinary anesthesiologist checked the physical status of the animal. This was followed by premedication with buprenorphine (0.01 mg/kg BW intramuscular) and medetomidine (0.07-0.1 mg/kg BW intramuscular). After 20 minutes the animal was transported to the CT-room, where the health status and degree of sedation were re-assessed.

Under sterile preconditioning a 14 G, 3.5-inch intravenous catheter was inserted into the left jugular vein and fixed to the skin. A fixed dose of midazolam (0.1 mg/kg BW) and ketamine (3mg/kg BW), with a variable dose of propofol (0.3-1.5 mg/kg BW) were administered intravenously to induce an anesthetic state. Then, under visual control using a laryngoscope, lidocaine 10% spray was used to desensitize the larynx. Following an allowance of 20-30 seconds, the animal's tracheas were intubated using an 11- or 12-mm ID endotracheal tube with the appropriate length. Consecutively, the animal was positioned in sternal recumbency with its front- and hind legs flexed forward. This body position was maintained for the duration of scanning.

On the CT table, the animals were immediately connected to a 22mm ID coaxial circle breathing system where artificial ventilation was maintained at a rate of 6-10 breaths per minute and 10 mL/kg BW of tidal volume. Connection of a pulse oximeter probe to the animal allowed for assessment and ensured adequate oxygenation, isoflurane was provided in oxygen at 0.8-2.0 Vol % throughout the whole scanning procedure.

Upon completion of the CT-protocol the breathing system was disconnected, and the animal was transferred to the operating room. Following reconnection to a breathing system with the same settings, a urinary catheter (Foley, size 9) was inserted into the urinary bladder for outflow and urine production monitoring. Hereafter, the animal was positioned for surgery, in sternal recumbency, on a flexible vacuum mattress for stable positioning. Monitoring included further ECG, SpO2, pharyngeal temperature, invasively measured arterial blood pressures (systolic, mean and diastolic) from a 20G cannula inserted into the auricular artery, inspired and expired oxygen, carbon dioxide and isoflurane tensions and spirometric monitoring of the adequacy of ventilation to the target EtCO2 of 35-40 mmHg.

Upon completion of the surgery, the animal was transported to the MR-room as fast as possible. Prior to the first BOLD-baseline sequence, a dose of 0.5 mg/kg BW rocuronium was administered as well as when the degree of muscle relaxation seemed insufficient for the completion of the procedure.

During all phases of the procedure, anesthesia was maintained, and when necessary, regulated by adjusting inspired isoflurane concentration and/or injection of additional propofol intravenously.

**CT examination**

CT examination of the sheep head and neck region was performed with a CT scanner (Philips Brilliance 16) and the following scanning parameters: slice thickness 1.5mm, increment 0.75, kV120, mAs 280, pitch 0.692, matrix 512x512, high resolution, collimation 16x0.75, rotation time 1s, sharp filter (c). The CT scan was used for neuro-navigation during the surgical procedure.

**Surgical procedures**

Following sternal recumbency onto a flexible vacuum mattress, a sterile surgical field was established by disinfection of the skin and placement of sterile draping. Neurosurgical navigation (Medtronic, S8 stealth station, Brainlab) was used to position an external ventricular drain (DePuys Synthes, Oberdorf, Switzerland) in the left lateral ventricle through an 11 mm burr hole.

**Euthanasia**

As soon as all planned data acquisition was terminated, the animal was euthanized under anesthesia by administration of pentobarbital (150 mg/kg BW) intravenously. Death was confirmed by transthoracic auscultation and readout of the attached monitoring instruments.
